# Telencephalic eversion in embryos and early larvae of four teleost species

**DOI:** 10.1101/2024.01.27.577540

**Authors:** Mónica Folgueira, Jonathan D. W. Clarke

**Author notes:** Corresponding author: Mónica Folgueira.

## Abstract

The telencephalon of ray-finned fishes undergoes eversion, which is very different to the evagination that occurs in most other vertebrates. Ventricle morphogenesis is key in order to build an everted telencephalon. Thus, here we use the apical marker *zona occludens 1* (ZO1) to understand ventricle morphology, extension of the *tela choroidea* and the eversion process during early telencephalon development of four teleost species: giant danio (*Devario aequipinnatus),* blind cavefish *(Astyanax mexicanus),* medaka *(Oryzias latipes)* and paradise fish (*Macroposus opercularis).* In addition, by using immunohistochemistry against tubulin and calcium binding proteins, we analyse the general morphology of the telencephalon, showing changes in the location and extension of the olfactory bulb and other telencephalic regions from 2 to 5 days of development. We also analyse the impact of abnormal eye and telencephalon morphogenesis on eversion, showing that *cyclops* mutants do undergo eversion despite very dramatic abnormal eye morphology. We discuss how the formation of the telencephalic ventricle in teleost fish, with its characteristic shape, is crucial event during eversion.

## 2. Introduction

Brain morphology is very variable in vertebrates, this diversity being specially manifested in the telencephalon (Nieuwenhuys et al., 1998; Northcutt, 2008; Aboitiz and Montiel, 2019; Briscoe and Ragsdale, 2019). These number of telencephalic forms can be classified into two main classes (Fig. 1): 1) ***everted telencephali***, present in actinopterygii (or ray-finned fishes) and partially in the sarcopterygii coelacanth (see Nieuwenhuys, 1965); and 2) ***evaginated telencephali,*** present in the rest of vertebrates (Holmgren, 1922; Johnston, 1911; Braford, 1995, 2009; Nieuwenhuys, 2009a, 2011). Despite the high variety in telencephalon morphology, the expression of molecules involved in patterning is highly conserved (Wilson and Rubenstein, 2000; Wullimann and Mueller, 2004; Briscoe and Ragsdale, 2019). This indicates that various developmental mechanisms, some of them still to be discovered, operate to generate remarkably different telencephalon morphologies under similar patterning signals.

**Figure 1.**
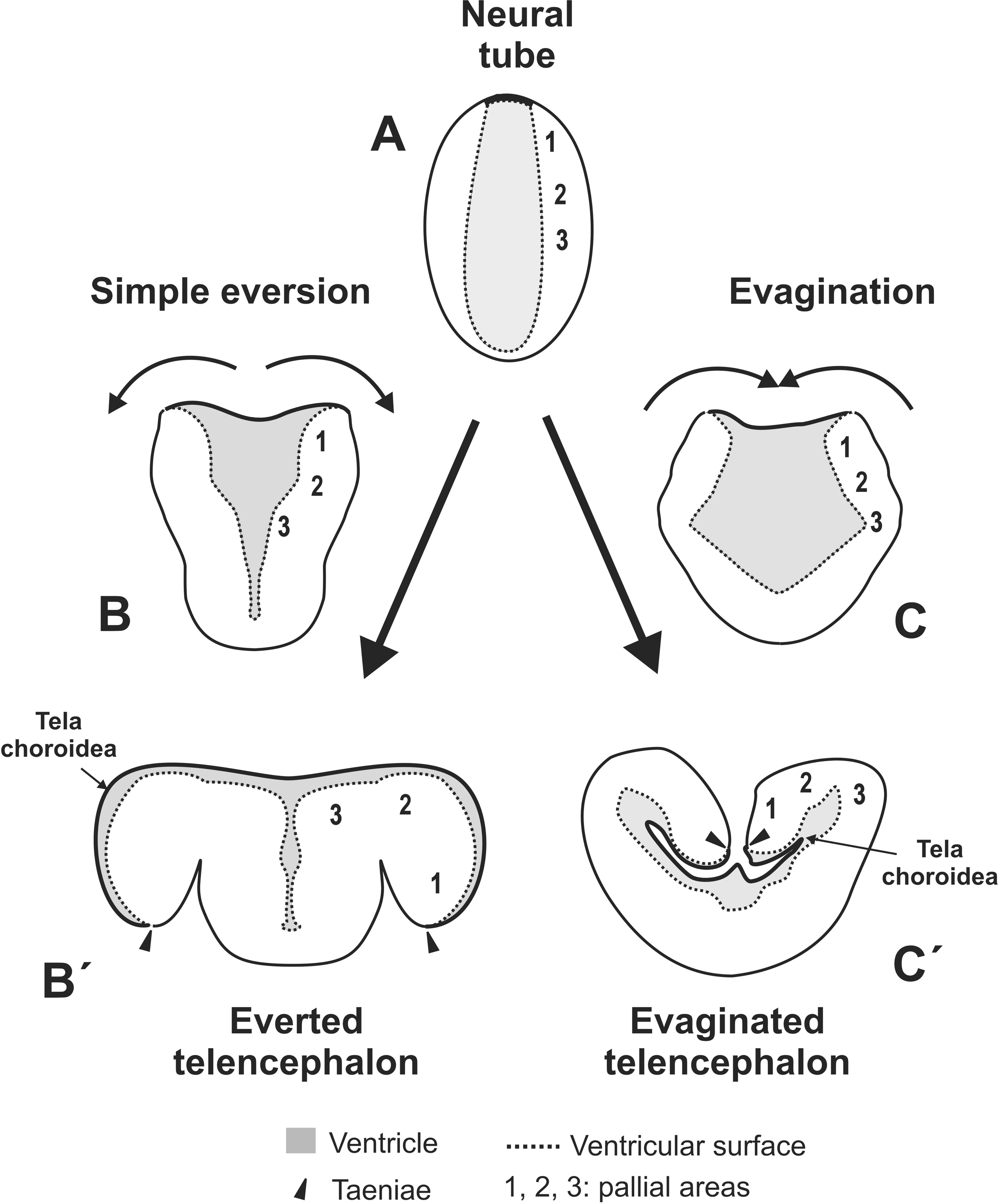
Schematic depicting the “classic eversion theory”. A-B’: eversion process, by which the neural tube (A) undergoes a morphogenetic process (B) that involves out-folding of its dorsal portion to result in an everted telencephalon in ray-finned fishes (B’). A-C’: evagination process, involving in-folding of the neural tube (C), resulting in an evaginated telencephalon (C’). Note the differences in the shape of the ventricle (grey) and location of the *choroid tela/plexus* (arrowheads) between everted and evaginated telencephali (B’ and C’). Asterisks mark the location of the *taeniae*. 1-3: pallial areas. Diagrams inspired in Gage drawings in plate viii (see Gage, 1893).

The everted telencephalon of actinopterygii (or ray-finned fishes) has a T-shaped ventricle that extends medially and laterally (Fig. 1B’), over the dorsal surface of the pallium, this being very different to the situation in evaginated telencephali (Fig. 1C’) (Braford, 1995, 2009; Nieuwenhuys, 2009a, 2011). Thus, in actinopterygii, a thin epithelial-like tissue, the *tela choroidea,* covers the dorsal surface of the telencephalon (Fig. 1B’). The degree of eversion varies between different species, so the *taeniae* (Fig. 1C’), that is, the places of attachment of the *tela choroidea,* are good landmarks for determining the degree of eversion (Nieuwenhuys, 1962, 2009b, 2011).

The eversion theory, proposed by Dr. Susanna Phelps Gage and Dr. František Karel Studničk, has been generally accepted to explain the development of the everted telencephalon. Based on the comparative analysis of brains from adult specimens of bowfin and newt, Dr. Gage proposed that the everted telencephalon is the result of an out-folding of the dorsal portion of the neural tube (Fig. 1A-B’; see also plate viii in Gage, 1893), which contrasts with the in-folding that happens during evagination in the rest of vertebrates (Fig. 1A-C’). Very soon after, Dr. Studnička proposed a very similar theory (Studnička, 1894, 1896; see Nieuwenhuys 2009a for a review). This eversion model has been referred to as “simple eversion theory” by Nieuwenhuys (2009a), and we will use this terminology from here onwards.

The main problem with the simple eversion theory is that it does not predict correctly the location of the pallial areas in the adult everted telencephalon, this being especially relevant for the location of the posterior zone of the pallium or Dp (Buttler, 2000; Wullimann and Mueller, 2004; Yamamoto et al., 2007; Nieuwenhuys, 2009a, 2011; Mueller et al., 2011). Thus, various modifications of the simple eversion theory have emerged trying to explain the location of the pallial areas in the adult telencephalon (Buttler, 2000; see Nieuwenhuys, 2009a for a review). Wullimann and Mueller (2004; see also Mueller et al., 2011) propose “a partial eversion model”, by which the pallial region that gives rise to Dp would not participate in eversion. Yamamoto and coworkers (2007), after analysing the location Dp and other telencephalic areas in adult goldfish, suggested that eversion not only involves a medio-lateral out-folding as proposed in the simple eversion model, but also a caudo-lateral one (see Fig. 4 in Yamamoto et al., 2007). After detailed analysis of the development of the zebrafish telencephalon, we proposed the “two-step model for telencephalic eversion” (Folgueira et al., 2012). Our results show that there are two critical events during early zebrafish development that results in an everted telencephalon by 5dpf (Fig. 2). First step is the shaping of the telencephalic ventricle from 18 to 24 hours post-fertilization (hpf), which includes generation of a lateral recess of the ventricle and medial to lateral rearrangements at the caudal end of the pallium (Fig. 2A-B, B’). This event results in the formation of the anterior intraencephalic sulcus (AIS), which is continuous with the optic recess ventrally (Affaticati et al., 2015). Second step is a rostro-caudal expansion of the telencephalon from 2 to 5 days post-fertilization (dpf) (Fig. 2C-D, C’-D’). Our results show that zebrafish telencephalon is everted by 5dpf (Fig. 2D’’; Folgueira et al., 2012), although further refinement happens from 5dpf into adult stages (Turner et al., 2022).

**Figure 2.**
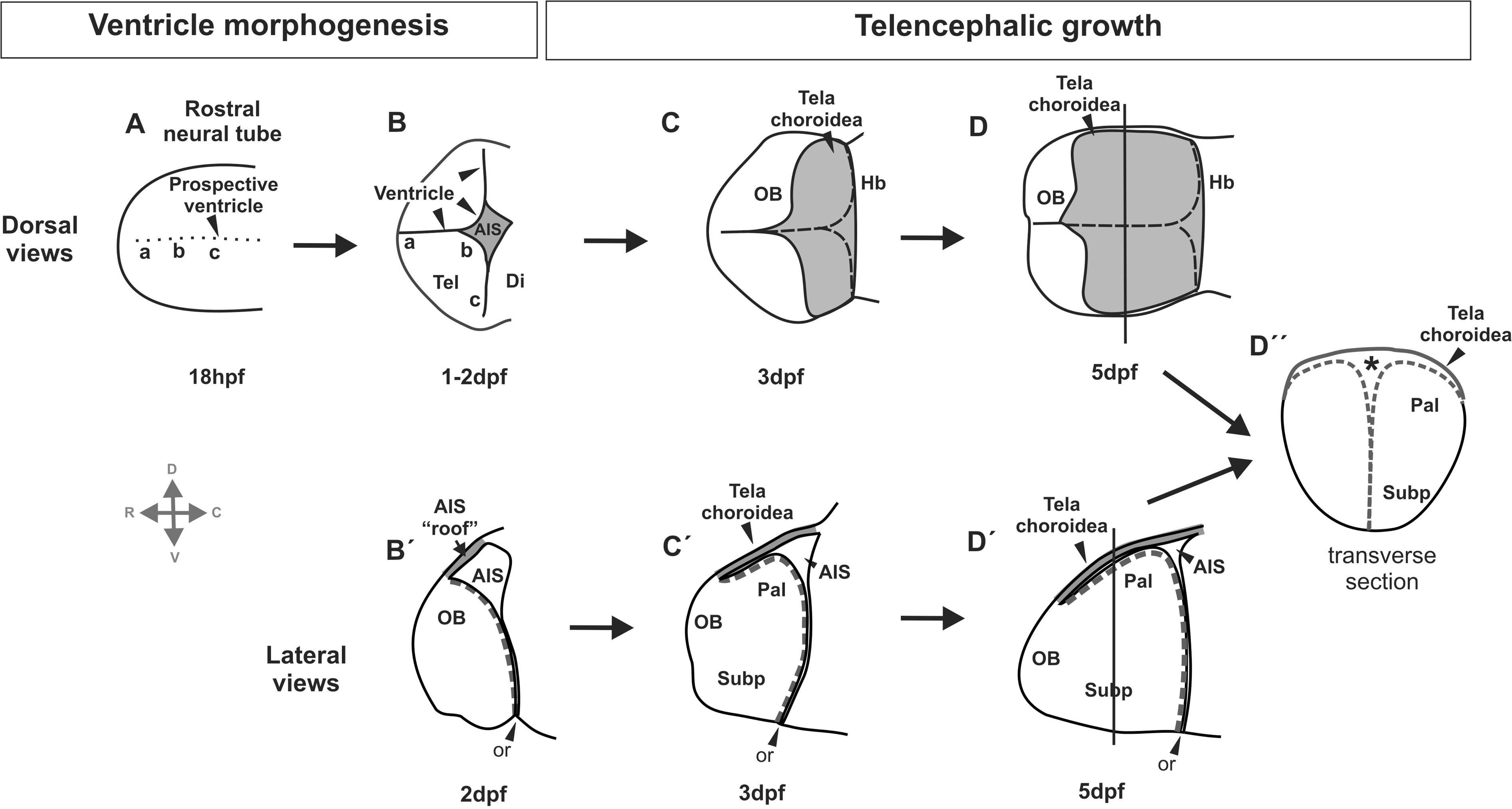
Schematic depicting the two-step model for telencephalic eversion (based on Folgueira et al., 2012). First is the formation of the telencepalic ventricle (A-B: dorsal views, B’: lateral view), including the AIS (marked in B, B’), leading to the medio-lateral rearrangement of neuroepithelial cells at the caudal end of the telencephalon (compare the positions of areas “a, b, c” in A and B). Next is the telencephalic expansion along the rostro-caudal axis from 48hpf to 5dpf (C-D in dorsal views, C’-D’ in lateral views). The roof of the AIS (B and B’) later becomes the *tela choroidea* (C-D and C’-D’). Dotted line in C-D marks the medial and posterior telencephalic ventricular surfaces. Dotted line in B’-D’ marks the dorsal and posterior telencephalic ventricular surfaces. D’’: transverse section through 5dpf telencephalon, showing that it is everted at this stage. Asterisk: ventricle. Abbreviations: AIS, anterior intraencephalic sulcus; Di: diencephalon; Hb, habenula, OB, olfactory bulb; or, optic recess; Pal, pallium; Subp, subpallium.

Given that key events of eversion happen during early development, it is surprising that most studies have analysed eversion mainly in the adult (Nieuwenhuys, 2009a, 2011), while very few studied developmental stages (Droogleever Fortuyn, 1961; Vázquez et al., 2002; Wullimann and Puelles, 1999; Striedter and Northcutt, 2006). This may be because of technical challenges when working with very small embryos and larvae, as well as difficulties to access the right developmental stages. In our previous study (Folgueira et al., 2012), we did analyse embryonic stages in detail, but only in one teleost species (zebrafish, *Danio rerio*). Of course, given the large number of species in the actinopterygii group, we cannot extrapolate that the two developmental events we observed in zebrafish are general to the group. Thus, rooting from our previous work in zebrafish, here we analyse telencephalon development in 4 freshwater tropical teleost species that have similar developmental speed to zebrafish: giant danio (*Devario aequipinnatus*), Pachon cavefish (*Astyanax mexicanus*), medaka (*Oryzias latipes*) and paradise fish (*Macroposus opercularis*).

All of them are members of the Euteleostei subdivision (Fig. 3) but show different degrees of telencephalic eversion in the adult (Fig. 4). As ventricle development is key for building an everted telencephalon, we studied the morphology of the telencephalic ventricle at early stages of development using the apical marker *zona occludens 1* (ZO-1). We also analysed changes in telencephalon morphology, with special attention to the development of the olfactory bulbs, pallium and fiber tracts. Finally, in order to test the robustness of the eversion process in presence of genetic perturbations that affect eye and telencephalon development, we analysed the morphology of the ventricle by 2pdf in *cyclops* zebrafish mutants, as well as the extension of the telencephalic ventricular surface by 5dpf. Our study draws a picture in which eversion is a robust process that has shared characteristics between all species studied, despite adult variability.

**Figure 3.**
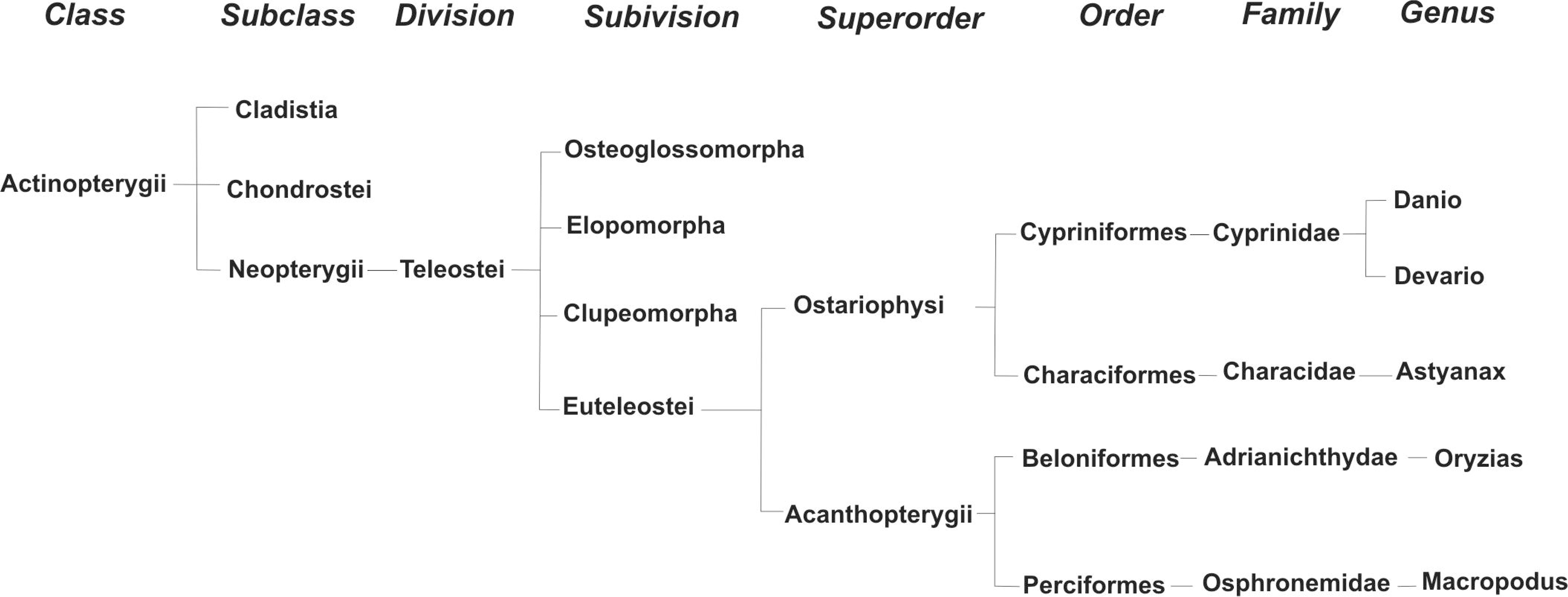
Phylogenetic tree showing the relationship and taxonomy of the species used in the present study (based in Nelson, 2006).

**Figure 4.**
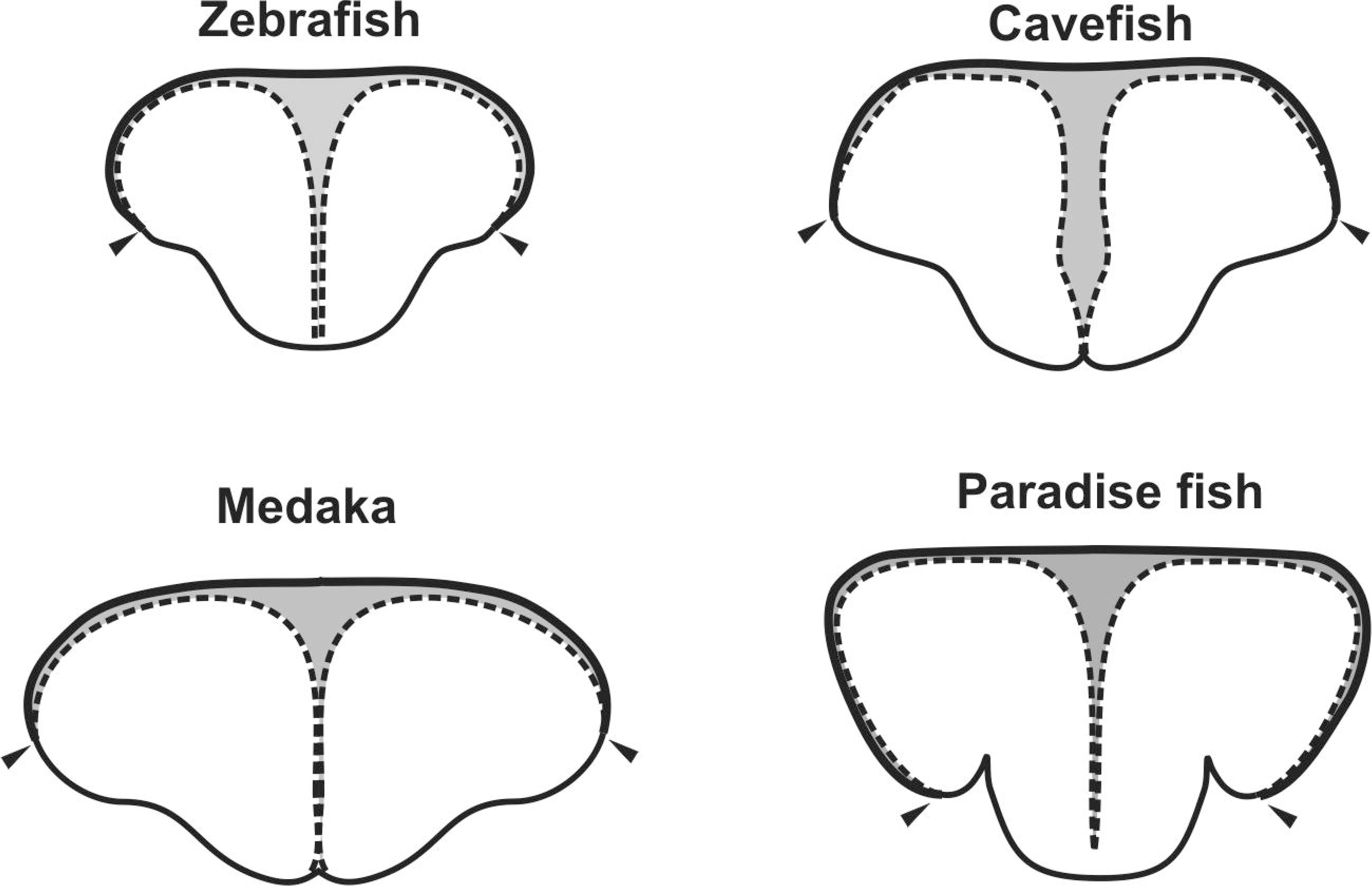
Schematic drawings showing the general morphology of transverse sections of the adult telencephalon of zebrafish, cavefish, medaka and paradise fish. The *taeniae* or points of attachment of the *tela choroidea* (arrowheads) allow visualizing the extension of the dorsal ventricular surface, that is, the degree of eversion. Dotted line: ventricular surface. Grey area: ventricle.

## 3. Material and methods

The fish species used in this study were (phylogeny following Nelson, 2006; Fig. 3): 1) **Zebrafish** : *Danio rerio, Fam. Cyprinidae, Superorder Ostariophysi, Euteleostei*; 2) **Giant Danio** : *Devario aequipinnatus, Fam. Cyprinidae, Superorder Ostariophysi, Euteleostei;* 3) **Cavefish (Pachon)** : *Astyanax mexicanus, Fam. Characidae, Superorder Ostariophysi, Euteleostei* ; 4) **Medaka** : *Oryzias latipes, Fam. Adrianichthyidae, Oder Beloniformes, Series Atherinomorpha, Superorder Acanthopterygii, Euteleostei* ; 5) **Paradise fish** : *Macroposus opercularis, Fam. Osphronemidae, Order Perciformes; Series Percomorpha, Superorder Acanthopterygii, Euteleostei*.

In addition, a zebrafish line carrying a mutation in *Nodal-related protein 2 (ndr2* ^m294^) was also used (Stemple et al., 1996; Solnica-Krezel et al., 1996; Sampath et al., 1998).

Individuals were obtained from various Fish Facilities, either collected directly by the authors or kindly donated by collaborators (see acknowledgements). Embryos and larvae were fixed in 4% paraformaldehyde (4% PFA) with 3% sucrose and then transferred to phosphate buffer saline 0.1 M pH 7.4 (PBS).

For the general morphology of the adult telencephalon shown in the introduction, transverse sections of cavefish and paradise fish stained with cressyl violet from our own collection were used. For medaka, schema is drawn based on Figure 1III in Okuyama et al. (2013). In this case, the location of the attachments of the *tc* or *taeniae* (the extension of the dorsal ventricular surface), was deduced based on the anatomy of the section. As no data was available for Giant danio, its close relative zebrafish was used.

All experiments were performed following the European Union Directive 2010/63/EU and Spanish Real Decreto 53/2013.

### Immunohistochemistry in embryos and larvae

Embryos and larvae used in this study were immunostained in whole mount following standard protocols (Folgueira et al., 2012; Turner et al., 2014, 2022). In brief, fish were transferred to methanol 100% and kept at -20°C for at least an hour. After rinsing in PBS with 0,5% Triton X100 (PBS-T), fish were treated with proteinase K, post-fixed in PFA and washed in PBS-T. Next, they were incubated in 10% normal goat serum in PBS-T for one hour and in the primary antibody or antibodies overnight (for dilutions see below). The next day, fish were washed in PBS-T and then incubated overnight with the secondary antibody or antibodies. Fish were then washed in PBS-T and, in the case of paradise fish and giant danio, transferred to 50% glycerol. Fish were mounted for imaging in either 2% agarose in PBS (zebrafish, cavefish and medaka) or in 1,2% agarose in 50% glycerol (giant danio and paradise fish).

Primary antibodies used were: mouse anti-*zona occludens 1* (ZO1, ThermoFisher-Invitrogen, 33-9100, dilution 1:300), mouse anti-acetylated α-tubulin (Sigma; Cat. T7451, dilution 1:250), rabbit anti-calretinin (CR, Swant; Cat. 7697; dilution 1:1000) and rabbit anti-calbindin D-28 (CB, Swant, CB38, dilution 1:1000). ZO-1 allows discerning the full extension of the telencephalic ventricle and *tela choroidea* (Grupp et al., 2010; Folgueira et al., 2012; Turner et al., 2022). Acetylated α-tubulin allows visualizing tracts and bundles (Chitnis and Kuwasa, 1990; Wilson et al., 1990, Ishikawa et al., 2004; Folgueira et al., 2012; Turner et al., 2022). Calcium binding proteins calretinin and calbindin D-28 label olfactory glomeruli (Castro et al., 2006; Germanà et al., 2007; Gayoso et al., 2011; Kress et al., 2015).

Secondary antibodies and dilutions used were: goat anti-rabbit Alexa Fluor 488 (Invitrogen, Cat. A-11034dilution 1:1000) and goat anti-mouse Alexa Fluor 568 (Invitrogen, Cat. A-11031, dilution 1:1000).

Fish were imaged using a Laser Scanning Confocal Microscope. For each stage and staining, an average of ten specimens were stained and at least two were imaged. Confocal stacks were processed using Fiji software (Schindelin et al., 2012).

Based on the neuromeric model, the anteriormost part of the brain is the optic chiasm (Puelles and Rubenstein, 1993; Wullimann, 2022). For simplicity, here we refer to “rostral” (towards the nose), “caudal” (towards the tail), “dorsal” and “ventral” for the description of results (see Herget et al., 2014; Porter and Mueller, 2020).

## 4. Results

### Telencephalon and ventricle morphology by 2 and 5 days of development

We performed whole-mount immunohistochemistry against the apical marker *zona occludens 1* (ZO-1) in giant danio, cavefish (Pachon), medaka and paradise fish in order to visualize the full extension of the ventricle, ventricular zone and *tela choroidea (tc)*.

In order to examine the development of the dorsal extent of the ventricular surface, we viewed ZO1 staining from dorsal. Giant danio, cavefish and paradise fish show similar telencephalon and ventricle morphology by 2dpf (Fig. 5A-B, A’-B’, D, D’; supplementary movie 1). The ventricle is very narrow at the rostral portion of the telencephalon, between both telencephalic lobes, but enlarges at the caudal end in the AIS. From dorsal view, the telencephalic surface shows a convex shape in all three species. In medaka, by 2dpf (about stage 28 of Iwamatsu, 2004) the ventricle is slightly different in shape and has a bigger cavity (Fig. 5C, C’; supplementary movie 2), which is similar to ventricle shape in 1dpf zebrafish development (not shown). In medaka, by 2dpf the ventricle does not show a convex shape dorsally, as it does in the other species, but it does more ventrally (Fig. 3C inset). Note that in all specimens, the ventricle is covered by the prospective *tela choroidea* (“roof” of the AIS; not shown).

**Figure 5.**
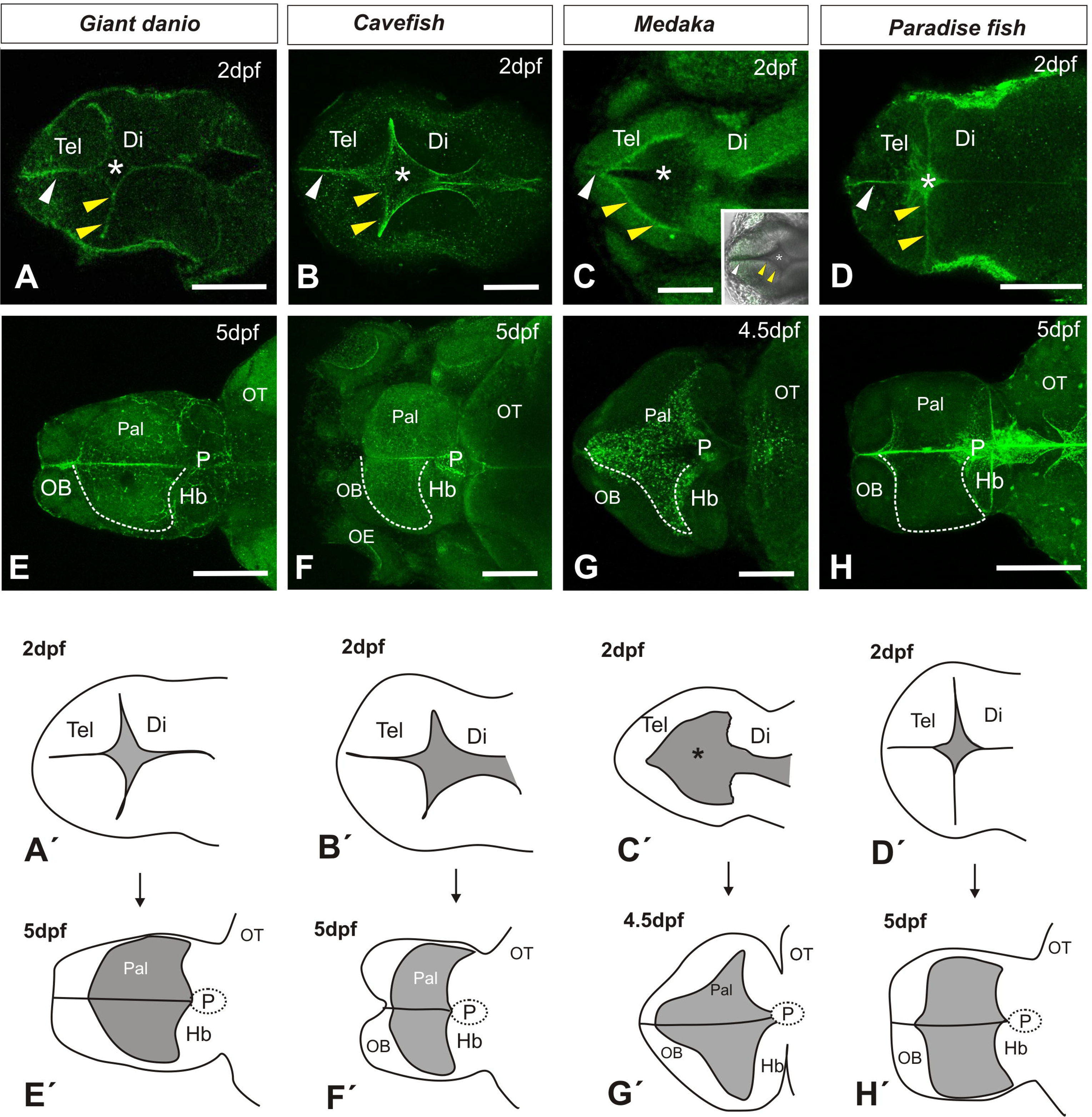
Dorsal views of the brain immunostained against ZO1 in 2dpf (A-D), 4.5dpf (G) and 5dpf (E-F, H) for the species studied (common name at the top). Schematics at the bottom of the figure (A’-H’) represent the general shape and main areas of the brain. A-D: Shape and extension of telencephalic-diencephalic ventricle. White arrowhead: narrow ventricular space between telencephalic lobes. Yellow arrowheads: posterior telencephalic ventricular surface towards the AIS. Asterisk: AIS. E-H: White dashed line in one side of the telencephalon marks the dorsal extension of the ventricular surface. A, B and D are confocal single scans. C, E-H: confocal stack projections. All are dissected brains, except F. A’-H’: marked in grey the dorsal roof of the AIS (2dpf), later becoming the *tela choroidea* (4,5-5dpf). Rostral to the left in all images. Abbreviations: Di, diencephalon; Hb, habenula, OB, olfactory bulb; OT: optic tectum; P, pineal; Pal, pallium; Tel, telencephalon. Scale bars: 100 µm in A-B, D-F, H; 50 µm in C and G.

Next we analysed the morphology of the telencephalic ventricle by 4/5dpf, in order to test whether there is a rostro-caudal expansion of the telencephalon that results in ventricular surface covering the dorsal telencephalon. By 4/5 days of development, the shape of the telencephalic ventricle has changed remarkably in the four species analysed (Fig. 5E-H, E’-H’; supplementary movies 3 and 4). Giant danio’s telencephalon by 5dpf (Fig. 5E, E’) largely resembles that of zebrafish (see Folgueira et al., 2012), being longer than in the other three species. Cavefish telencephalon is slightly shorter along the A-P axis (Fig. 5F, F’), but wider than giant danio. Medaka telencephalon ventricle has a triangular shape by 4.5dpf (stage 33-34 of Iwamatsu, 2004; Fig. 5G, G’) that contrasts with the telencephalon shape in the other three species. In this species, it is noteworthy the thickening of the telencephalic walls that has happened from 2dpf to 5dpf. Finally, paradise fish’s telencephalon is similar to that of giant danio (Fig. 5H, H’), but slightly shorter and wider rostrally. In all species, as a consequence of telencephalon growth, ventricular surface and *tc* now covers the dorsal surface of the telencephalon (Fig. 5E-H, E’-H’; movies 5-8; note that ZO-1 does not allow distinguishing ventricular zone *vs*. *tc*). Our results show that, as previously observed in zebrafish, it is a rostro-caudal expansion of the telencephalon that results in the ventricular surface reaching the dorsal telencephalic surface.

In order to analyse in more detail the rostro-caudal elongation phase of telencephalon development, we examined the expression of α-tubulin in giant danio, paradise fish and cavefish in lateral view. This marker allows observation of major neuropil fibers and bundles (Chitnis and Kuwada, 1990; Wilson et al., 1990), among other structures. As the ventricle and ventricular surface cannot be clearly visualized with this marker, we used the olfactory bulb (telencephalon) and habenulae (diencephalon) as landmarks. To confirm the location of the olfactory bulb, in addition we studied the expression of the calcium-binding proteins calretinin (in giant danio and paradise fish) and calbindin D-28 (in cavefish), which label a subset of olfactory glomeruli (see Castro et al., 2006; Gayoso et al., 2011; Kress et al., 2015).

In giant danio, the olfactory bulb glomeruli are clearly visible, being located close to the habenulae by 2dpf (Fig. 6A). Calretinin staining labels a subset of 2 glomeruli in the olfactory bulb (Fig. 6A, inset). By 5dpf, there is great enlargement of the telencephalon (Fig. 6B). The olfactory bulb is now located much farther away from the habenulae, consequence of the growth of the telencephalic lobes. At this stage, calretinin labelling is more extensive, labelling at least 5 olfactory glomeruli (Fig. 6B inset). The staining does not allow delineation of the exact limit of the different subdivisions of the telencephalic lobes. However, it is clear that there is a rostro-caudal expansion between the olfactory bulb and the habenulae between 2dpf and 5dpf.

**Figure 6.**
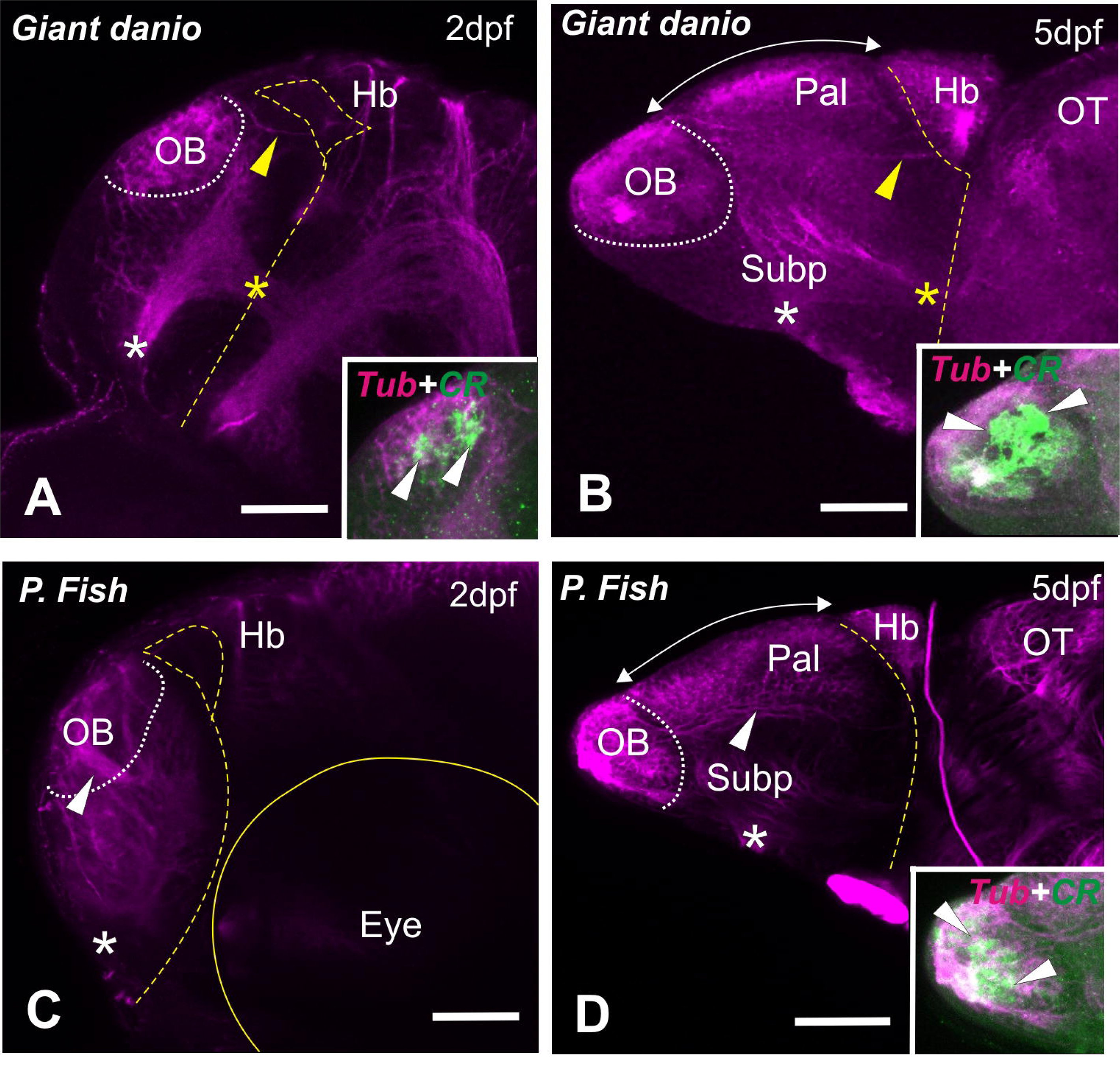
Tubulin immunostaining (Tub, magenta) in giant danio (top, A-B) and paradise fish (bottom, C-D) showing telencephalon morphology by 2dpf (A, C) and 5dpf (B, D). Insets in A and B show olfactory bulb glomeruli labelled with calretinin (CR, green) together with tubulin staining. White asterisk: anterior commissure. Yellow asterisk: forebrain bundle. Yellow arrowhead: *stria medullaris*. White arrowheads in insets: olfactory glomeruli. White arrowhead in C: olfactory nerve. White arrowhead in D: tract, maybe corresponding to lateral olfactory tract (see text). White dashed line marks the posterior and ventral limits of the olfactory bulb. Yellow dashed line marks AIS and optic recess. Yellow line in C: eye. Rostral to the left. Abbreviations: CR, calretinin; Hb, habenula; OB, olfactory bulb; OT, optic tectum; Pal, pallium; Subp, subpallium; Tub, alpha-tubulin. Scale bars: 50 µm.

In paradise fish, the olfactory nerve can be clearly observed as a thick axon bundle that reaches the olfactory bulb, which by 2dpf is also located near the habenulae (Fig. 6C). By 5dpf, there is also a dramatic expansion of the telencephalon, especially caudal to the olfactory bulbs (Fig. 6D), which are now placed further away from the habenulae. Again, there is a dramatic increase in domains in the telencephalic lobe with a clear rostro-caudal expansion.

In cavefish, we analysed acetylated α-tubulin expression by 2dpf, 3dpf and 5dpf (Fig. 7). By 2dpf and 3dpf, the olfactory bulb can be clearly discerned with tubulin staining (Fig. 7A-F); calbindin D-28 labels a single olfactory glomerulus and some olfactory cells (Fig. 7B-C, E-F). Again, by 2dpf and 3dpf the olfactory bulb is located quite close to the habenulae (Fig. 7A-F) By 5dpf the telencephalon shows a dramatic change in shape, being much more elongated than previous stages (Fig. 7G-I). Calbidin D-28 labels 4-5 glomeruli in the olfactory bulbs (Fig. 7H-I), which are now quite distant from the habenulae. Cavefish clearly shows an elongation of the telencephalon along the rostro-caudal axis from 2/3dpf to 5dpf.

**Figure 7.**
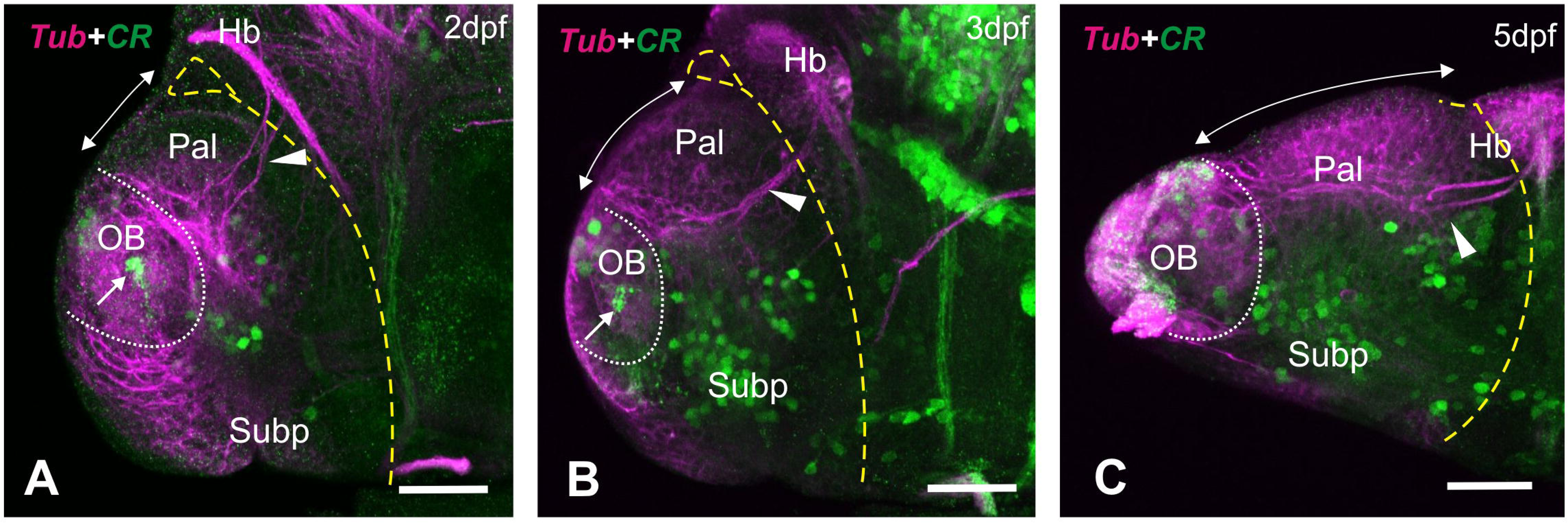
Tubulin and calretinin immunostaining of cavefish by 2dpf, 3dpf and 5dpf. A-: Lateral views of cavefish brains labelled against tubulin (magenta) and calretinin (green). Rostral to the left. White dashed line marks limits of the olfactory bulb. Yellow dashed line marks AIS and optic recess. Double headed arrow: extension of the dorsal ventricle. White arrowheads: *stria medullaris*. Yellow arrowhead: tract, maybe lateral olfactory tract. Abbreviations: CR, calretinin; Hb, habenula; OB, olfactory bulb; OT, optic tectum; Pal, pallium; Subp, subpallium; Tub, tubulin. Scale bars: 50 µm.

Detailed description of Calbindin D-28 expression is out of the scope of the manuscript, but it is noteworthy that our results show cell-specific expression in the central and peripheral nervous system. Briefly, we observed Calbidin D-28 immunoreactivity in cells of olfactory bulb, subpallium, thalamus, pretectum, optic tectum and in sensory neurons of the trigeminal ganglion by 2dpf (the trigeminal ganglion was not present in 3 and 5dpf dissected tissue).

### Does eversion occur with abnormal neurulation in Cyclops mutants?

One hypothesis for eversion implicates the small available space within the skull of ray-finned fishes, so the telencephalon must squeeze between the olfactory organs and the eyes (Ariëns Kappers, 1908; Striedter and Northcutt, 2006). If that were the case, abnormal neurulation that result in eye malformations might impact eversion in two ways: a) by altering the available space within the skull and b) by changing ventricle morphology, as the AIS is continuous with the optic recess ventrally. Thus, we analysed ventricle morphology in cyclops *ndr2* ^te262c^ -/- embryos, which show different degree of eye fusion or cyclopia, as well as variability in the position of the olfactory epithelium. In order to see how alteration in neurulation and eye morphogenesis impacts eversion in these embryos (England et al., 2006; Aquilina-Beck et al., 2007), we performed staining against ZO-1 at 2dpf and 5dpf. Our results show that the AIS in cyclops mutants show similar shape to that in wild-type by 2dpf, but with a smaller cavity (Fig. 8A, C). Ventrally, the optic recess does show important morphology differences between Cyclops and wild-types (supplementary movies 5 and 6), a consequence of eye malformation in the mutants. The AIS is also similar in shape to that of the other species analysed in this work (compare 8C with 5A-D). By 5dpf, the ventricle in cyclops mutants shows altered morphology compared to wild-types (Fig. 8B, D, supplementary movies 7 and 8). However, importantly the telencephalon has everted in cyclops, as the ventricular surface and roof of the AIS has expanded over the dorsal telencephalon as in wild-types. Thus, although showing differences in shape compared to wild-type fish, cyclops mutants develop an everted telencephalon despite alterations in neurulation and eye morphology.

**Figure 8.**
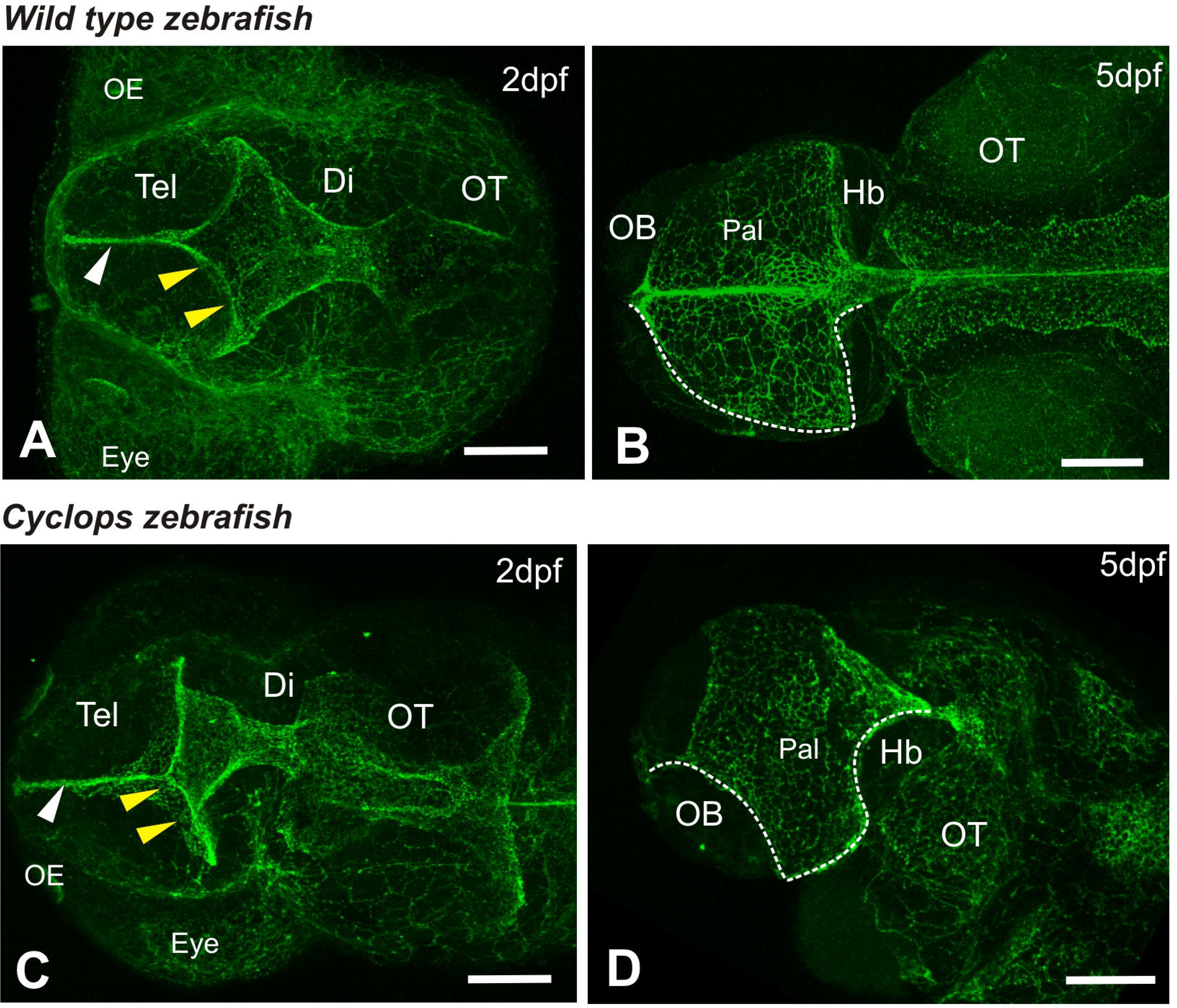
Dorsal view of 2dpf and 5dpf wild types (A-B) and Cyclops mutants (C-D) immunostained against ZO-1, showing the extension of the ventricle in the forebrain. White arrowhead points to ventricle between both telencephalic lobes. Yellow arrowheads point to telencephalic portion of the ventricle towards the AIS. White dashed line in B and D mark the full dorsal extension of the ventricle in one side of the brain. A and C are whole embryos. B and D is the dissected brain only. Rostral to the left. Abbreviations: Di, diencephalon; Hb, habenula, OB, olfactory bulb; OE: olfactory epithelium; Pal, pallium; Tel, telencephalon. Scale bars: 50 µm.

## 5. Discussion

### 5.1 An eversion model for teleosts

We examined telencephalon morphology of four teleost species in order to assess whether their development supported the “two-step model of eversion” we proposed earlier based on detailed analysis of zebrafish brain morphogenesis (Folgueira et al., 2012). The first step in our model involves the generation of the anterior intraencephalic sulcus (AIS) (Fig. 9), a deep ventricular recess between telencephalon and diencephalon (von Kupffer, 1896). This occurs soon after the formation of the midline ventricular seam at the end of the neural rod stage. The formation of the AIS generates a posterior ventricular surface at this stage and shifts medial ventricular progenitors to a more lateral location (Figs. 2A-B, 9A-B). The posterior ventricular surface curves like the outside surface of a balloon, resulting in its characteristic convex shape. Dorsally the AIS is covered by a thin ventricular roof called the *tela choroidea* (tc).

**Figure 9.**
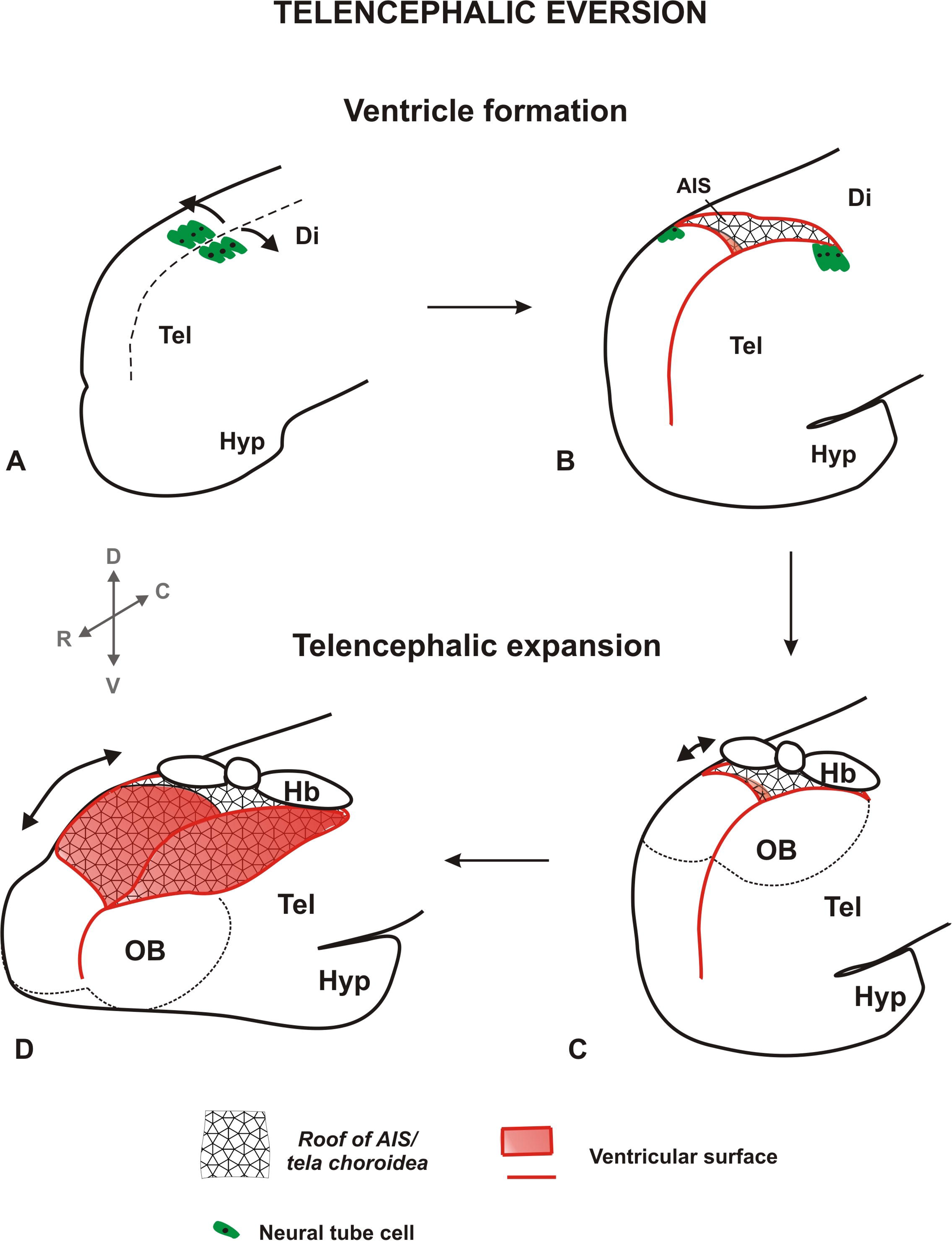
Three dimensional schematic of early telencephalic eversion in teleosts. AIS: anterior intraencephalic sulcus; Di: diencephalon; Hb, habenula; Hyp: hypothalamus; OB, olfactory bulb; Tel: telencephalon.

Our results reveal the morphology of the AIS in four species by 2 days post fertilization (Figure 5), showing the presence of a posterior ventricular space in all species, which results in the medio-lateral shift of ventricular cells in posterior telencephalon. The morphology of the AIS has also been shown by micro-CT in the rosy bitterling (*Rhodeus ocellatus*, Cypriniformes; Yi et al., 2022), being very similar to the morphology in the species we analysed. Given that some of these species are phylogenetically distant, these data suggest that the formation of the AIS, with its characteristic shape, is a common early key step in telencephalic morphogenesis in teleosts. There are small variations in the precise morphology of the AIS across the species analysed (most notably seen in medaka), which may be accounted for by variations in the speed of morphogenesis, rather than fundamental differences in tissue shaping. Striedter and Northcutt (2006) have previously analysed the shape of the telencephalic ventricle in horizontal sections of catfish. Remarkably, the first step of our model (Figs. 2A-B, 9A-B) is very similar to Striedter and Northcutt’s model presented in horizontal sections (compare Figure 7E-H in Striedter and Northcutt, 2006 with Fig. 2B). Taken together these observations confirm the formation of the AIS generates the convex arrangement of the teleostean ventricular surface and is an early key step in the telencephalic eversion process.

The second step in our model for eversion involves a major vector of telencephalic growth along the rostro-caudal axis. This greatly enlarges the distance between olfactory bulb rostrally and the habenulae caudally. During this phase, the posterior wall of the telencephalon bulges upwards, towards the thin roof of the AIS or prospective *tela choroidea*, creating the characteristic dorsal surface of the telencephalic ventricular surface and the narrow ventricular space between this and the *tela choroidea* (Folgueira et al., 2012; Figs. 2B’-D’, 9C-D). Here we show that a rostro-caudal expansion of the dorsal surface of the telencephalon occurs between days 2 and 5 of development in all four species examined, probably even starting by 1dpf. Similar results were also obtained in the rosy bitterling by Yi et al. (2022), showing an expansion of the ventricular surface of the AIS from 165 to 235hpf. Furthermore, our stainings for neuropil confirms the increasing distance between OB and habenulae during this period in giant danio, paradise fish and cavefish (unfortunately we were unable to get sufficient embryos to check this in medaka). These results are consistent with our proposal that the formation of the dorsal ventricular surface and the *tela choroidea* of the telencephalon in teleosts is principally formed by a rostro-caudal expansion of dorsal telencephalon and not by an out-folding of the neural tube originally proposed as the major morphogenetic process of the simple eversion theory (Gage, 1893, Studnička, 1894, 1896; and reviewed in Nieuwenhuys 2009a).

Our observations in embryos of zebrafish, giant danio, paradise fish, cavefish and medaka are consistent with the proposal that the two-step model of eversion operates across diverse teleost species, and argue against neural tube mediolateral out-folding being the principal morphogenetic mechanism of eversion. We did not observe any medial to lateral rearrangements from 2dpf to 5dpf in zebrafish (Folgueira et al., 2012), nor in the species examined here. We do observe medial to lateral rearrangements at the caudal end of the telencephalon as a consequence of ventricle formation (Figs. 2A-B, 9C-D), which will be discussed in detail later. We cannot rule out the possibility that mediolateral rearrangements beyond 5 days of development do contribute to the adult form of the telencephalon. Certainly the 5dpf telencephalon is different in morphology and size to that in the adult (personal observations), and important changes occur in the telencephalon from 5 to 20dpf (Turner et al., 2022). Thus it is very likely that cell proliferation and differentiation beyond 5dpf contributes to expansion of the everted surface, and may generate some later medial to lateral rearrangements (Mueller et al., 2011; Porter and Mueller, 2020; Turner et al., 2022).

We propose our two-step model of eversion is likely to be applicable to most, if not all, teleost species. In addition to zebrafish and the species analysed here, Striedter and Northcutt (2006) show similar morphology of the ventricle in catfish (*Ictalurus punctatus*, Fam. Ictaluridae, Order Siluriformes). Thus, the same morphology is shared by 6 teleost species. One could argue that this number of species is still too modest to confidently generalize the model to a whole group with more than 30,000 species. However, the fact that the same morphology is present in phylogenetically distant species supports the hypothesis of the model being valid to the whole group. We do wonder to what extend the model could be also applied to other groups of ray-finned fish, such as Cladistia and Chondrostei. We believe it could be valid to explain the appearance of the everted morphology of the telencephalon in these groups, but not to account for the appearance of the internal ventricle in the olfactory bulb. Interestingly, Tavighi and coworkers (2016) studied ventricle development in beluga sturgeon (*Huso huso*). By 1dpf, the telencephalic ventricle has very similar morphology to that observed here in teleosts (see Fig. 1 in Tavighi et al., 2016). However, by 6dpf ventricle morphology is much more complex, showing more grooves than in teleosts (see fig. 2 in Tavighi et al., 2016). Vazquez et al. (2002) studied telencephalon development in *Acipenser naccarii* but they do not show horizontal sections or dorsal views, so it is not possible to determine the whole morphology of the ventricle. Of special interest would be to analyse ventricle morphology during early stages of development in bichir (*Polypterus*). The pallium has a very characteristic shape in this species, being a rather thin bended wall (Reiner and Northcutt, 1992, Nieuwenhuys 2009b, 2011)). This is likely to be the consequence of the short migration of mature neurons from the ventricular surface during development. Our prediction would be that the ventricle in early stages of bichir development would resemble caudally that of teleosts, but this needs testing. Eversion may not be exclusive of actinopterygii, as the coelacanth (*Latimeria*, *Fam. Coelacanthidae*), the only surviving crossopterygian, has a partly everted telencephalon (see Nieuwenhuys, 1965). Of course it would be extremely interesting to analyse the development of this living fossil to understand eversion, but needless to say this is completely out of experimental reach.

Our results in cyclops (*ndr2* homozygous) mutants indicate that early eversion is a very robust process. These fish carry a mutation that affects nodal signalling, resulting in abnormal ventral patterning of the neural tube (Sampath et al., 1998). Although these embryos show different degrees of cyclopia, in all cases we observed the characteristic shape of the AIS by 2dpf and a fully everted telencephalon by 5dpf. Thus the AIS acquires its characteristic shape despite the mutation in nodal signalling or the alteration in midline patterning. This shape must be, at least partially, the result of changes in apico-basal cell shape, as a result of cytoskeleton rearrangements, as described for the midbrain-hindbrain ventricle (Gutzman et al. 2008; Lowery and Sive, 2009). Other mechanisms, such as differential cell proliferation and apoptosis might also be involved (Gutzman et al. 2008). There are a number of zebrafish mutants that show alteration in ventricle morphogenesis (Lowery et al., 2009), so it would be interesting to analyse how eversion is affected in these mutants.

### 5.2. How does eversion affect the location of cell populations in the pallium?

Our analyses show that in actinopterygians the early formation of the AIS at the telencephalic/diencephalic border invariably leads to medial to lateral repositioning of neuroepithelial cells at the caudal end of the telencephalon (Figs. 2A-B, 9A-B; as also implied (Striedter and Northcutt, 2006) or explicitly proposed in previous models (Yamamoto et al., 2007). However, this medio-lateral remodelling during early stages of development does not generate a fully everted telencephalon, as proposed by the classic eversion model. It is necessary further growth of the telencephalon to generate a fully everted telencephalon, with its characteristic dorsal ventricular zone and *tela choroidea*. Based on the anatomy of the telencephalon in goldfish, Yamamotós model proposes a medio-lateral folding at the caudal end of the telencephalon (Yamamoto, 2007). We believe this medial to lateral shift proposed by Yamamoto and coworkers (2007) happens, at least partially, during early ventricle morphogenesis, as a result of ventricle opening, as shown for other species (Folgueira et al., 2012; present results). This early medio-lateral morphogenesis of the posterior telencephalon generates the AIS in both the Yamamoto and our own model. This step helps explain the mature location of the posterior area of the pallium (known as Dp), but whose location is not easily explained by the simple medio-lateral out folding of the neural tube proposed in the classic eversion model (Nieuwenhuys 2009a, 2011).

It is hard to infer what consequences the medio-lateral rearrangements at the caudal end may have on the organization of all the caudal telencephalic areas. Fate map studies will be necessary to fully understand how the medio-lateral shift that happens during ventricle formation results in the organization of the telencephalic areas observed in the adult. This is even more important when results in zebrafish have shown that medial and lateral regions of the pallium have different developmental timing (Dirian et al., 2014), which inevitably will have an impact in the adult pallium arrangement. Previous studies trying to unravel telencephalon homologies tend to compare late stages of development or even adult stages of everted and evaginated telencephali. In order to compare and establish homologies between both modes of telencephalon organization it is necessary to look at early, easily comparable stages before analysing later ones. This will help not only to build the whole picture of how the different telencephalic areas acquire their final position, but also to establish homologies between the telencephalic regions.

### 5.3. Why does eversion occur?

It is not clear when, why and how everted telencephali emerged in actinopterygii. A recent study of the fossilized brain of the extinct ray-finned fish *Coccocephalus wildi* showed that this species seemed to have an evaginated brain (Figueroa et al., 2022), suggesting that an everted telencephalon might have evolved later than thought until now. Ariëns Kappers was, to our knowledge, first to suggest in 1908 that eversion could happen as a consequence of lack of space within the skull. Striedter and Northcutt (2006) recovered this theory and developed it further, relating the appearance of the everted telencephalon in actinopterygii with their reproductive strategy. These authors noted the small space within the skull, which correlates with body size in the embryo, independently of adult size. This would force embryonic telencephalon to use all the available space within the skull, undergoing eversion instead of evagination to minimise ventricular volume (Striedter and Northcutt, 2006). However, results in the extinct *Coccocephalus wildi* now challenges this theory, as it seems that this species had an evaginated telencephalon despite restricted available space within the skull (Figueroa et al., 2022).

Early ventricular morphology is a key aspect of telencephalon morphogenesis and could mark the divergence between generating an everted or evaginated telencephalon (Fig. 7; see also Aboitiz and Montiel, 2019). Neurulation, leading to the formation of a ventricle with a specific shape, could mark already the divergence that leads to generate an everted or evaginated telencephalon. In most models, a hollowed neural tube is the common starting point for everted and evaginated telencephali. However, this is an over simplification, as teleostean neurulation and lumen formation is different to other vertebrates (Harrington et al., 2009; Aboitiz and Montiel, 2019). It would be interesting to further study how changes in neurulation and ventricle size impacts eversion. This could be achieved, for instance, by mechanically removing the optic cups during development. Another way could be to use fish mutants with alterations in forebrain patterning. For instance, in zebrafish *rx3* homozygous mutants eye field cells acquires a telencephalic fate. These embryos have larger AIS (see Fig. 8 in Stigloher et al., 2006), which could result in an increased ventricular surface dorsally by 5dpf, hence a more everted telencephalon. However, this has not been tested. More studies are necessary to clarify why and how eversion appeared during evolution, and what effect experimental manipulation has on eversion, as discussed by Striedter and Northcutt (2006).

## 8. Data availability statement

The raw data used for the production of this article will be made available by the corresponding author, without undue reservation.

## 9. Author contribution

Work conceptualization and design: MF and JDWC; data acquisition and analysis: MF; original draft preparation: MF; review and editing of manuscript, MF and JDWC. Both authors have read and agreed to the final version of the manuscript.

## 10. Acknowledgements

MF would like to acknowledge past work colleagues at University College London (UCL) for kindly providing most of the biological material used in this study: Dr. Yoshiyuki Yamamoto (UCL) for cavefish; Dr. Mathias Carl (University of Trento, Italy) for medaka; Dr. Máté Varga (Eötvös Loránd University, Budapest, Hungary) for Paradise fish and Dr. Rodrigo Young (University College London) for Giant danio. MF would also like to acknowledge Dr. Steve Wilson (UCL) and members of his lab for helpful comments and discussion.

## 11. Conflict of interests

Author declares no conflict of interest

## 12. Funding

The authors received no financial support for this research.

## 13. Graphical Abstract Text

The telencephalon of ray-finned fishes undergoes eversion, not evagination as in the rest of vertebrates. We show that the formation of the ventricle followed by a rostro-caudal telencephalic expansion is key for eversion in teleosts.

## 14. Graphical Abstract Image

Figure 9

## 15. Research Highlights

The developmental events that lead to an everted telencephalon in ray-finned fishes are not well understood. Here we show that the morphology of the telencephalic ventricle is quite similar between 5 species of teleosts, being a key event during eversion. We also show the transformation that the telencephalon undergoes from 2 to 4/5 days of development, including growth that leads to an expansion along the rostro-caudal axis and that results in a fully everted telencephalon. Our results support the idea that these two key events, ventricle formation and telencephalon elongation, are at the core of the formation of an everted telencephalon in teleosts. In addition, we show zebrafish cyclops mutants develop an everted telencephalon despite very abnormal neurulation that alters ventricle and eye morphology. Our results provide experimental basis for the only model of eversion based in development data.

## Notes

### Competing Interest Statement

The authors have declared no competing interest.

